# Deletion of INPP5E in the murine retina impairs axoneme formation and prevents photoreceptor disc morphogenesis

**DOI:** 10.1101/2020.11.29.403030

**Authors:** Ali S. Sharif, Cecilia D. Gerstner, Martha A. Cady, Vadim Y. Arshavsky, Christina Mitchell, Guoxin Ying, Jeanne M. Frederick, Wolfgang Baehr

## Abstract

INPP5E (pharbin) is a ubiquitously-expressed, farnesylated phosphatidylinositol polyphosphate 5’-phosphatase which modulates the phosphoinositide composition of membranes. INPP5E resides in primary cilia, and mutations or loss of INPP5E are associated with ciliary dysfunction. INPP5E missense mutations of the phosphatase catalytic domain cause Joubert syndrome in humans, a syndromic ciliopathy affecting multiple tissues including brain, liver, kidney and retina. We show that, differing from other primary cilia, INPP5E is present in the wildtype photoreceptor inner segment and absent in the outer segment--a modified primary cilium dedicated to phototransduction. We generated *Inpp5e*^F/F^;Six3Cre (in short, ^ret^*Inpp5e^-/-^*) mice which exhibit a rapidly progressing rod-cone degeneration nearly completed by postnatal day 21 (P21) in the central retina. Mutant cone outer segments contain vesicles instead of discs as early as P8. While P10 mutant outer segments contain phototransduction and structural proteins, they do not form axonemes and fail to elaborate disc membranes. Connecting cilia of ^ret^*Inpp5e^-/-^* rods appear normal, although IFT-B/A particles accumulate at their distal ends suggesting disrupted intraflagellar transport. These results show that ablation of INPP5E does not impair the secretory pathway responsible for delivery of outer segment-specific proteins, but blocks axonemal extension and prevents disc morphogenesis.

---

INPP5E is a farnesylated phosphatidylinositol 5’-phosphatase (1,2) catalyzing the hydrolysis of the 5’ phosphate from PI(4,5)P_2_ (PIP2), and PI(3,4,5)P_3_ (PIP3) (reviewed in (3,4)). Phosphatidylinositol polyphosphates (PIPs) play major roles in cell division, integral membrane protein transport, vesicular trafficking, and ciliary formation and function (4-6). INPP5E is associated with Joubert (JS) and MORM syndromes (7,8) caused by missense mutations in the phosphatase domain. JS is a syndromic ciliopathy affecting the brain, eyes, kidney and liver (9,10) presenting with ataxia, hyperpnea, polydactyly, molar tooth sign in brain and retinitis pigmentosa or Leber congenital amaurosis.

Ten phosphatidylinositol 5’-phosphatases (INPP5A, B, D, E, F, G, H, J, K and INPPL1) are present in mammals, and of these, three (A, B, E) are farnesylated (11). Germline mouse knockouts of INPP5E and INPP5K in mice are embryonically lethal suggesting non-redundant roles for some phosphatase isoforms in various cells or sub-compartments (7,9,12). *Inpp5e^-/-^* mice (deletion of exons 7 and 8) died soon after birth; E18.6 *Inpp5e^-/-^* embryos showed developmental arrest at the optic vesicle stage before appearance of the optic cup. *Inpp5e^-/-^* E18.5 embryos displayed multiple cysts, polydactyly and skeletal abnormalities (7). *Inpp5e^-/-^* embryos were anophthalmic suggesting severe consequences of INPP5E deletion during early eye development (7). A second *Inpp5e^-/-^* mouse model (deletion of exons 2-6) confirmed the JS phenotype and identified disordered sonic hedgehog-dependent patterning during embryonic development (12). Conditional deletion of INPP5E in kidneys resulted in severe polycystic kidney disease (PKD) and hyperactivation of PI3K/Akt and mTORC1 signaling (13). Similarly, deletion of the Inpp5e gene in zebrafish led to cystic kidneys (14).

Using transfection of IMCD3 cells with plasmids encoding FLAG- or EGFP-INPP5E, INPP5E localized predominantly to primary cilia (15,16). As a C-terminally farnesylated protein, INPP5E is chaperoned to the cilium by the prenyl-binding protein PDEδ (16-18). Ciliary localization in hTERT-RPE1, 293T and IMCD3 cells was confirmed using a polyclonal anti-INPP5E antibody (14,19-21). *Inpp5e^-/-^* mouse embryonic fibroblasts (MEFs) developed primary cilia suggesting INPP5E 5’-phosphatase activity is not essential for ciliogenesis but mutant cilia were more sensitive to resorption during the cell cycle (5,6,9). In contrast to primary cilia, EGFP-INPP5E introduced to rod photoreceptors by neonatal electroporation distributed to inner segments (IS) (22). INPP5E immunolabeling of dissociated rods similarly showed a prominent IS signal and an additional non-uniform staining of the outer segment (OS) (23).

In this study, we deleted INPP5E from the retina by mating *Inpp5e*^F/F^ mice (12) with Six3Cre transgenic mice. Using multiple techniques, we show that INPP5E localizes to the IS of wildtype (WT) photoreceptors. *^ret^Inpp5e^-/-^* ROS start to degenerate by P10. While P10 connecting cilia of mutant rods are nearly normal in length, shortened ROS are devoid of recognizable discs. IFT-A and IFT-B particles accumulate in the proximal *^ret^Inpp5e^-/-^* OS, suggesting defective intraflagellar transport (IFT). Thus, deletion of INPP5E primarily impairs axoneme extension and disc morphogenesis of both rods and cones.

## Results

### Generation of retina-specific *Inpp5e* knockout mice

INPP5E is a 72 kDa protein carrying a proline-rich domain in the N-terminal region, a large phosphatase active site encoded by exons 3-9, a coiled-coil domain and a CAAX motif for C-terminal farnesylation (**Fig. 1A**). Mutations in human INPP5E associated with JS are located predominantly in the phosphatase domain. The mouse *Inpp5e* gene consists of ten exons, producing two splice variants (**Fig. 1B**). The full-length variant is predicted to be farnesylated and anchored to membranes. The shorter variant lacking exon 10 and the CAAX box is predicted to be soluble.

**Figure 1.**
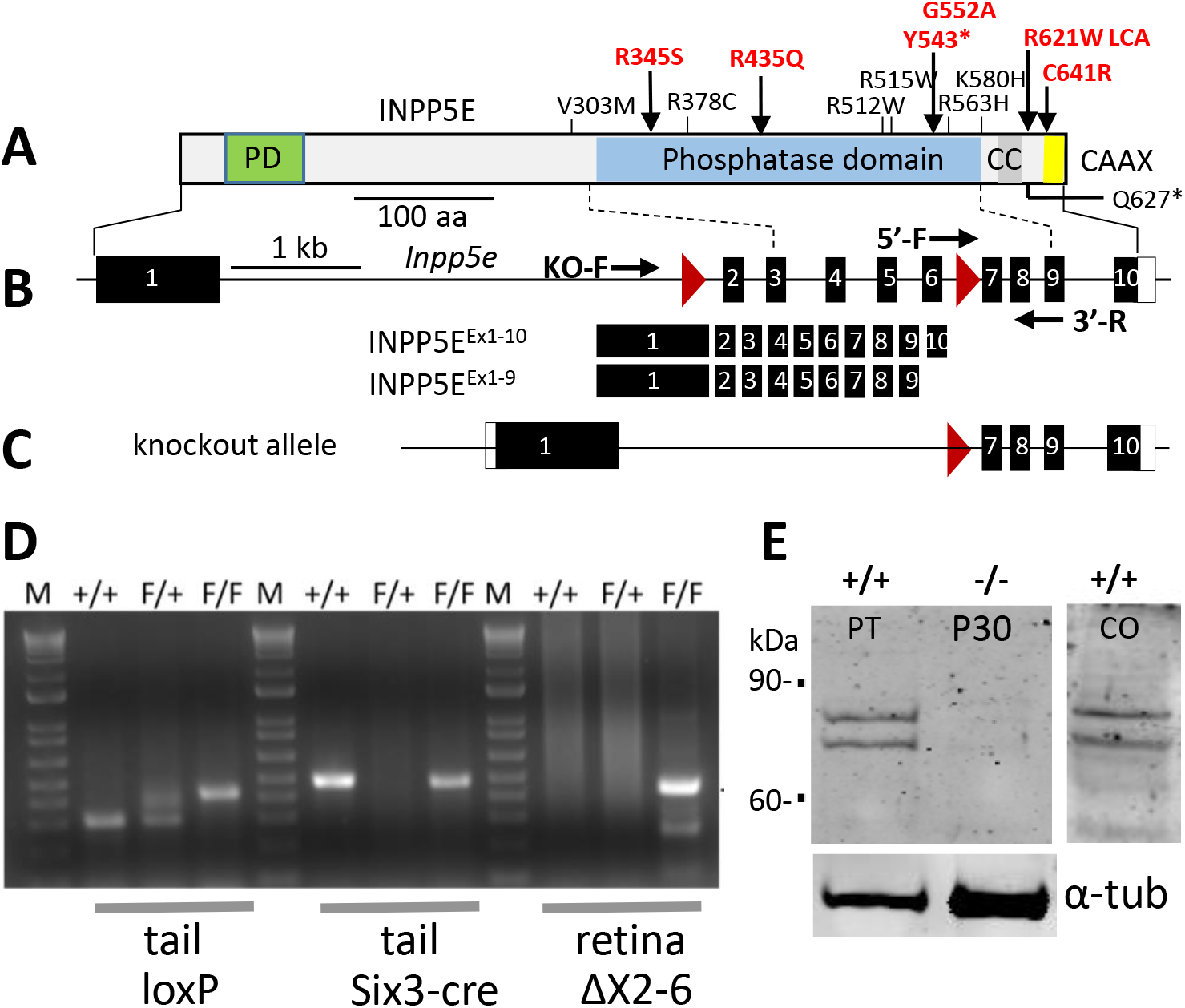
Retina-specific knockout of INPP5E. **A,** INPP5E major functional domains. PD, proline-rich domain; CC, coiled-coil domain; CAAX, motif for prenylation. Approximate positions of missense mutations associated with Joubert syndrome are indicated; mutations shown in red display a retina phenotype. **B,** the murine *Inpp5e* gene with 10 exons (black bars) and two LoxP sites (red triangles). Mouse *Inpp5e* gene expresses two splice variants, both isoforms contain the phosphatase domain. The shorter variant lacks the last exon carrying the CAAX motif. **C,** knockout allele in which exons 2-6 are deleted. **D,** PCR genotyping protocol. Assay for WT and floxed alleles (left), for absence or presence of Six3Cre (middle), and for deletion of exons 2-6 (right). Amplicons of lanes 4, 8 and 12 indicate a knockout mouse. **E**, INPP5E immunoblot with commercially available antibody (ProteinTech) (at P30) and custom antibody (Covance) (at P21), revealing the presence of two variants INPP5E^Ex1-10^ and INPP5E^Ex1-9^ in the mouse retina. Both variants were deleted in the knockout.

To generate retina-specific knockout (^ret^*Inpp5e^-/-^*) mice, *Inpp5e*^F/F^ mice were bred with Six3Cre mice (24). LoxP sites placed in introns 1 and 6 specify deletion of most of the phosphatase domain (**Fig. 1C**). LoxP, Six3Cre and knockout genotyping are shown (**Fig. 1D**). Immunoblotting with two independent antibodies against human INPP5E (ProteinTech) and a mouse N-terminal fragment of INPP5E (custom-made at Covance, Inc.) confirmed the presence of two splice variants in WT retinal lysates and deletion of both isoforms in the conditional knockout line (**Fig. 1E**).

### Electroretinography

We recorded whole-field scotopic and photopic electroretinograms (ERGs) at P15 and P21 at light intensities ranging from - 1.63 to 2.38 log cd s/m^2^ (**Fig. 2)**. Scotopic a-waves (**Fig. 2A, D** and scotopic b-waves (**Fig. 2B, E**) of knockout mice were barely recordable at P15 and P21. Scotopic a-waves of heterozygous littermates displayed normal a-wave amplitudes at P21 (**Fig. 2D)**, indistinguishable from WT suggesting haplosufficiency. Surprisingly, the P15 ^ret^*Inpp5e^-/-^* photopic ERG b-wave amplitudes were still recordable (**Fig. 2**) presumably owing in part to a slower degeneration of cones in the peripheral retina. At P21 ^ret^*Inpp5e^-/-^* photopic ERG b-wave amplitudes were much reduced (**Fig. 2F**). The heterozygous b-wave amplitudes were somewhat reduced, but the reduction was not statistically significant.

**FIGURE 2.**
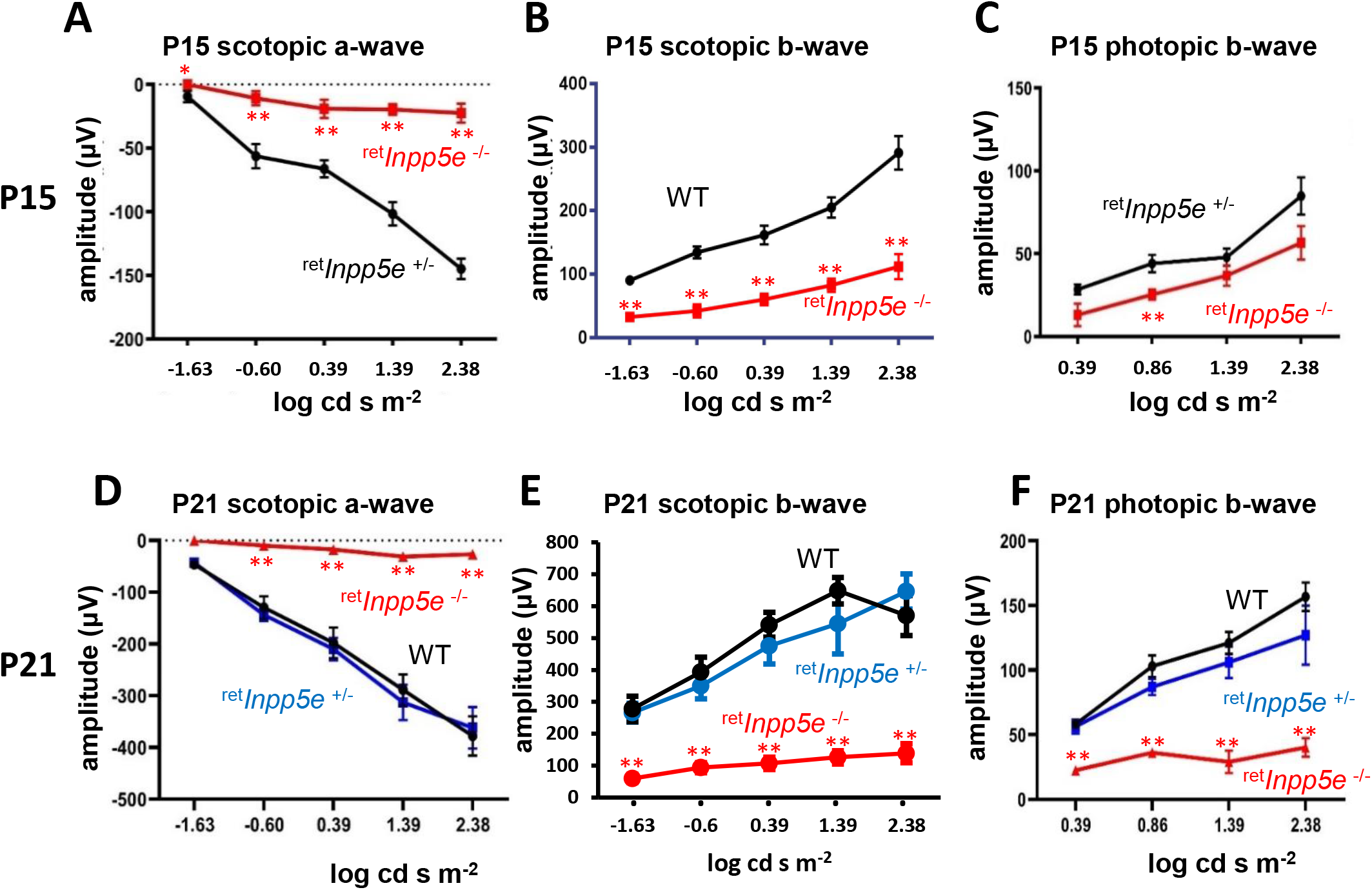
Electroretinography of control and ^ret^*Inpp5e^-/-^* mice. **A-F,** scotopic and photopic ERG measurements conducted at P15 (A-C) and P21 (D-F). A-wave and b-wave amplitudes were plotted as a function of flash intensity. Note significant photopic b-wave activity of the ^ret^*Inpp5e^-/-^* retina at P15 (C) and minor haploinsufficiency at P21 (F). P15 control n=7 and KO n=6. P21 control n=3 and KO n=4. Data analyzed via *t*-Test: two samples assuming equal variance. Data are presented as mean ±SEM. All a-wave scotopic amplitudes are less than \*\**p*<0.01 and all b-wave amplitudes are less than \**p*<0.05. Mutant mice were compared with their respective WT and heterozygous littermates. P15 control n=7 and KO n=6. P21 WT n=4, Hets n=3, and KO n=4. There is no statistical significance at P15 photopic b-wave amplitude at 1.38 and 2.38 log cd s m^-2^.

### ^ret^*Inpp5e^-/-^* photoreceptors undergo rapid degeneration

Plastic sections of ^ret^*Inpp5e^-/-^* and littermate WT control retinas collected at P10, P15 and P21 (**Fig. 3A-F**) confirmed rapid photoreceptor degeneration. At P10, ^ret^*Inpp5e^-/-^* retinas (**Fig. 3D**) showed only slightly reduced ONL thickness compared to WT (**Fig. 3A**), but the distance between RPE and ONL was substantially reduced suggesting abnormal size of IS and/or OS. At P15 nuclear tiers were reduced by ~30% (**Fig. 3E**) and at P21 only one row of nuclei, presumably cones, was present (**Fig. 3F**), consistent with rapid *rd1*-like degeneration. Evaluation of the ONL thickness across the entire retina (**Fig. 3G-I**) showed a 40% reduction in the central retina at P15 and nearly complete obliteration of the central ONL at P21. The less affected ONL thickness at the retinal periphery (**Fig. 3I**) is consistent with lower Cre expression in peripheral retina under the Six3 promoter (24,25). The ^ret^*Inpp5e^-/-^* inner retina, including the INL and GC layers, was practically unaffected, as judged by near normal morphology of P10, P15 and P21 cryosections (**Fig. 3D-E**).

**FIGURE 3.**
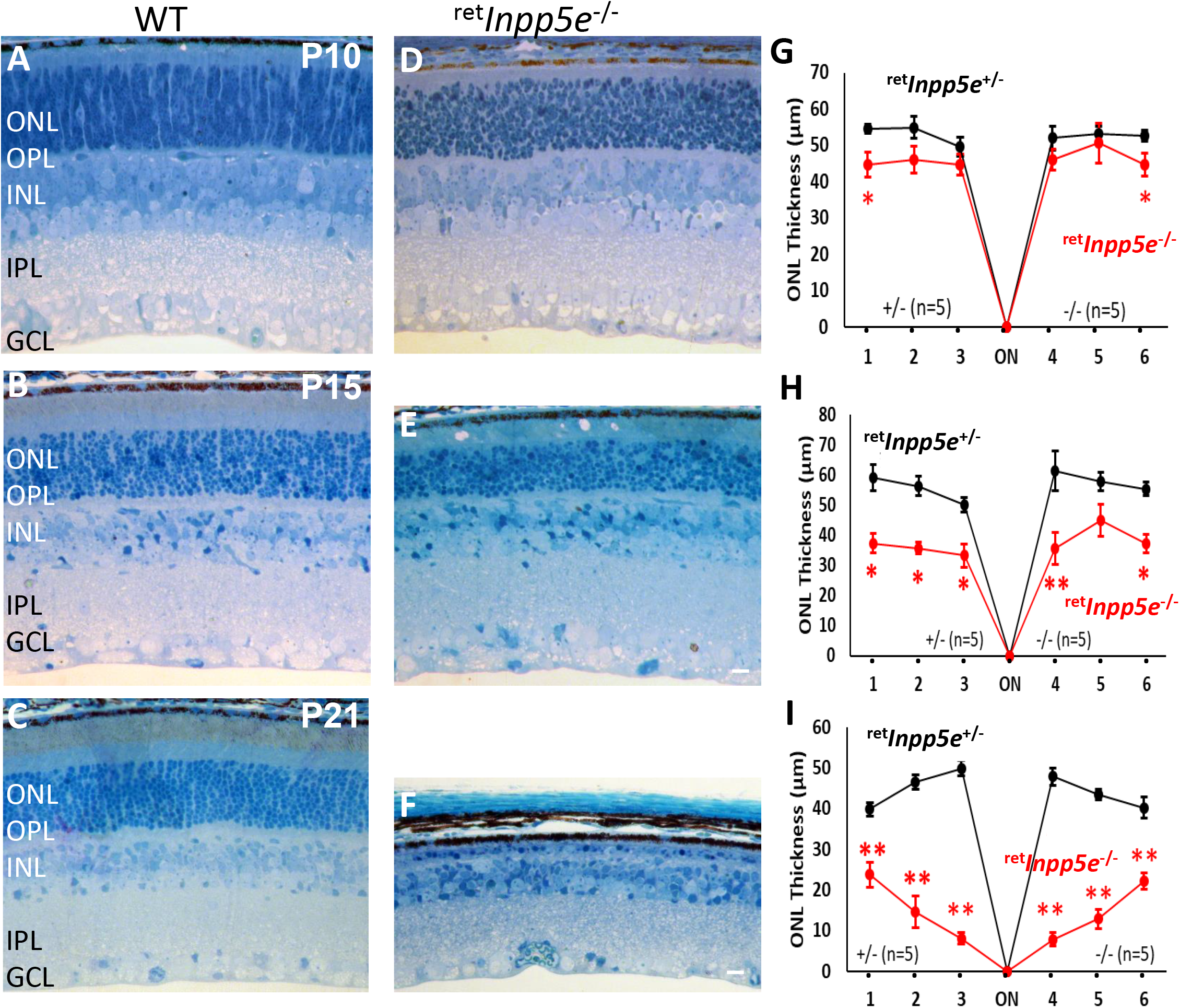
Progressive photoreceptor degeneration in ^ret^*Inpp5e^-/-^* mice. **A-F**, plastic sections obtained from the central retinas of WT (A-C) and ^ret^*Inpp5e^--/-^* (D-F) mice at P10, P15 and P21. The ONL thickness in knockout mice is nearly normal at P10, but is reduced to 7-8 rows of nuclei at P15. Only one nuclear row remains in the knockout ONL at P21. *Scale bar*, 10 μm. **G-I**, ONL thickness of WT versus knockout retinas, measured at 500 μm intervals from the optic nerve head (ON) at P10 (G), P15 (H) and P21 (I). Note that, at P21, the *^ret^Inpp5e^-/-^* ONL is preserved more at the periphery. Five control and knockout animals were analyzed at each time point. Data are presented as mean ±SEM. n =5; student *t*-test \**p*_0.05; \*\**p*_0.01.

### INPP5E is a photoreceptor inner segment protein

Immunolocalization of INPP5E in photoreceptors was performed with a well-characterized anti-INPP5E antibody “PT” (ProteinTech) raised against a His-tagged fusion protein corresponding to the N-terminal half of human INPP5E. WT cryosections analyzed at P15 (**Fig. 4A**) and P21 (**Fig. 4C**) revealed the presence of INPP5E in IS with traces in the ONL, and absence in OS. In knockout retinas, the IS content of INPP5e was significantly reduced at P15 (**Fig. 4B**) and undetectable at P21 (**Fig. 4D**).

**FIGURE 4.**
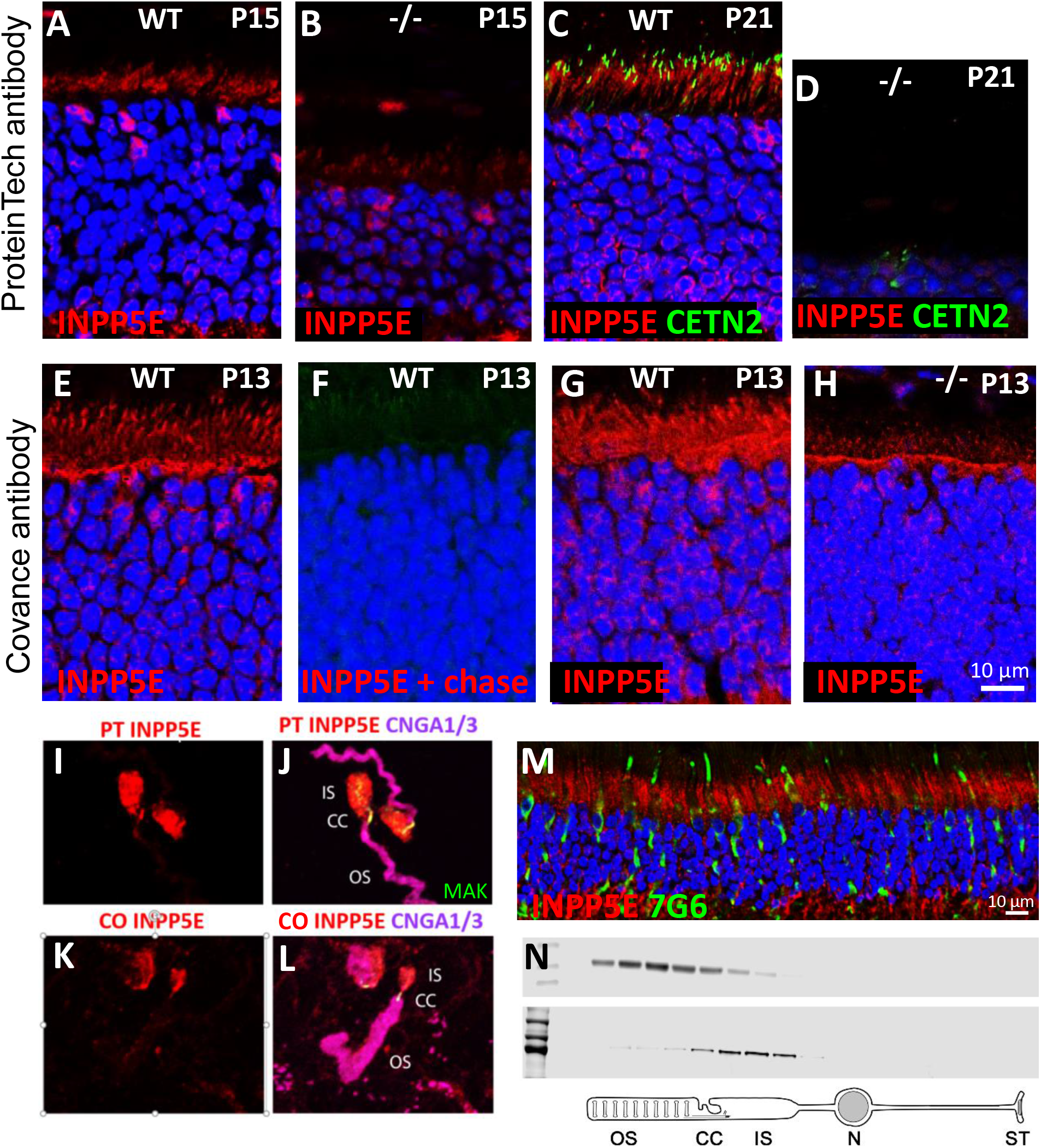
INPP5E localization in photoreceptors. **A-D,** INPP5E immunostaining in WT (A, C) and ^ret^*Inpp5e^-/-^* (B, D) retinal cryosections using the ProteinTech antibody at P15 (A, B) and P21 (C, D). In ^ret^*Inpp5e^-/-^* mice, INPP5E is weakly detectable at P15 and completely absent at P21. **E-H,** analysis of the INPP5E custom-made Covance antibody specificity in WT (E-G) and knockout retina cryosections (H) at P13. In F, Covance antibody was preabsorbed with the mouse N-terminal peptide thereby obliterating immunoreactivity. **I-L,** INPP5E distribution in isolated, lightly fixed mouse rods using the ProteinTech (I, J) and Covance (K, L) antibodies (red). Outer segments are labeled by an antibody against CNGA1/3 (magenta) and CC by an antibody against MAK (male germ cell associated kinase, green). **M,** INPP5E immunostaining in postmortem human macula from a 71 year-old donor. INPP5E (red) is detected robustly in photoreceptor IS. Cones are identified by an antibody recognizing primate cone arrestin (mAb 7G6, green), nuclei are stained by DAPI (blue). **N**, tangential sectioning of a rat retina in 5 μm sections. Proteins in individual sections were probed with an anti-rhodopsin (top) and anti-INPP5E (PT, bottom) antibodies. Below, a cartoon depicting approximate locations of individual sections.

To verify the specificity of the human antibody in the mouse, we generated an independent antibody “CO” against the N-terminal peptide (MPSKSASLRHTEAC) of mouse INPP5E. As its sequence is distinct from the human homolog, the antibody is predicted to be mouse-specific. This antibody showed strong immunoreactivity in the entire IS, along the outer limiting membrane (OLM), with traces in the ONL (**Fig. 4E**). The immunoreactivity was erased when the antibody was saturated with the antigenic peptide (**Fig. 4F**). Anti-INPP5E immunoreactivity was still detectable in knockout sections at P15 with the PT antibody (**Fig. 4B**) and at P13 with CO antibody (**Fig. 4H**) suggesting ineffective clearing of INPP5E or non-specific binding. Differences in IS antigen localization using the PT antibody (fixed sections, **Fig. 4A**) and CO antibody (slightly fixed sections, **Fig. 4E**) are most likely caused different fixation protocols (see *Experimental Procedures*).

Dissociated mouse rods (23) were probed with both antibodies (**Fig. 4I-L**). OS were identified by anti-CNGA1/3 and the connecting cilium (CC) by anti MAK (male germ cell associated kinase) antibodies. These results clearly showed absence of INPP5E in the OS and its prominent presence in both IS and CC.

The INPP5E localization was further confirmed in macula of a 71 year-old human postmortem donor where INPP5E was detected in the IS of rods and cones (**Fig. 4M**). This pattern differs from INPP5E localization in the primary cilium where INPP5E is delivered with the aid from the solubilization factor PDEδ (PDE6D) and the cargo displacement factor ARL3-GTP (16).

### Serial tangential sectioning of rat retinas

To verify the IS localization of INPP5E beyond any doubt, we complemented immunolocalization analysis with the technique combining serial tangential sectioning of the retina and immunoblotting (26). A western blot of proteins from serial sections cut through a WT rat retina starting from the photoreceptor tips was probed with antibodies against rhodopsin (the OS marker) and INPP5E (**Fig. 4N**). This analysis revealed the presence of a single INPP5E band in sections corresponding to IS (the rat *Inpp5e* gene expresses no splice variants) and only trace protein amounts in the sections representing OS.

### Absence of PDEδ does not affect INPP5E localization

The IS location of INPP5E was unchanged in cryosections obtained from *Arl3^-/-^* mice (22), suggesting that it is not delivered in a complex with the prenyl-binding protein PDEδ. We verified that cellular distribution of farnesylated INPP5E is not dependent on PDEδ by analyzing its localization patterns in *Pde6d^-/-^* photoreceptors (**Fig. 5**). OS of WT cones contain geranylgeranylated PDE6α’ (**Fig. 5A**) and farnesylated GRK1 (**Fig. 5B**) whose transport to cone OS is completely impaired in the absence of PDEδ (**Fig. 5D, E**). In contrast, the localization of farnesylated INPP5E was insensitive to absence of PDEδ (**Fig. 5C, F**). It is conceivable that the small amount of INPP5E detected at the CC (**Fig. 4J, L)** is trafficked there by complexing with PDEδ.

**FIGURE 5.**
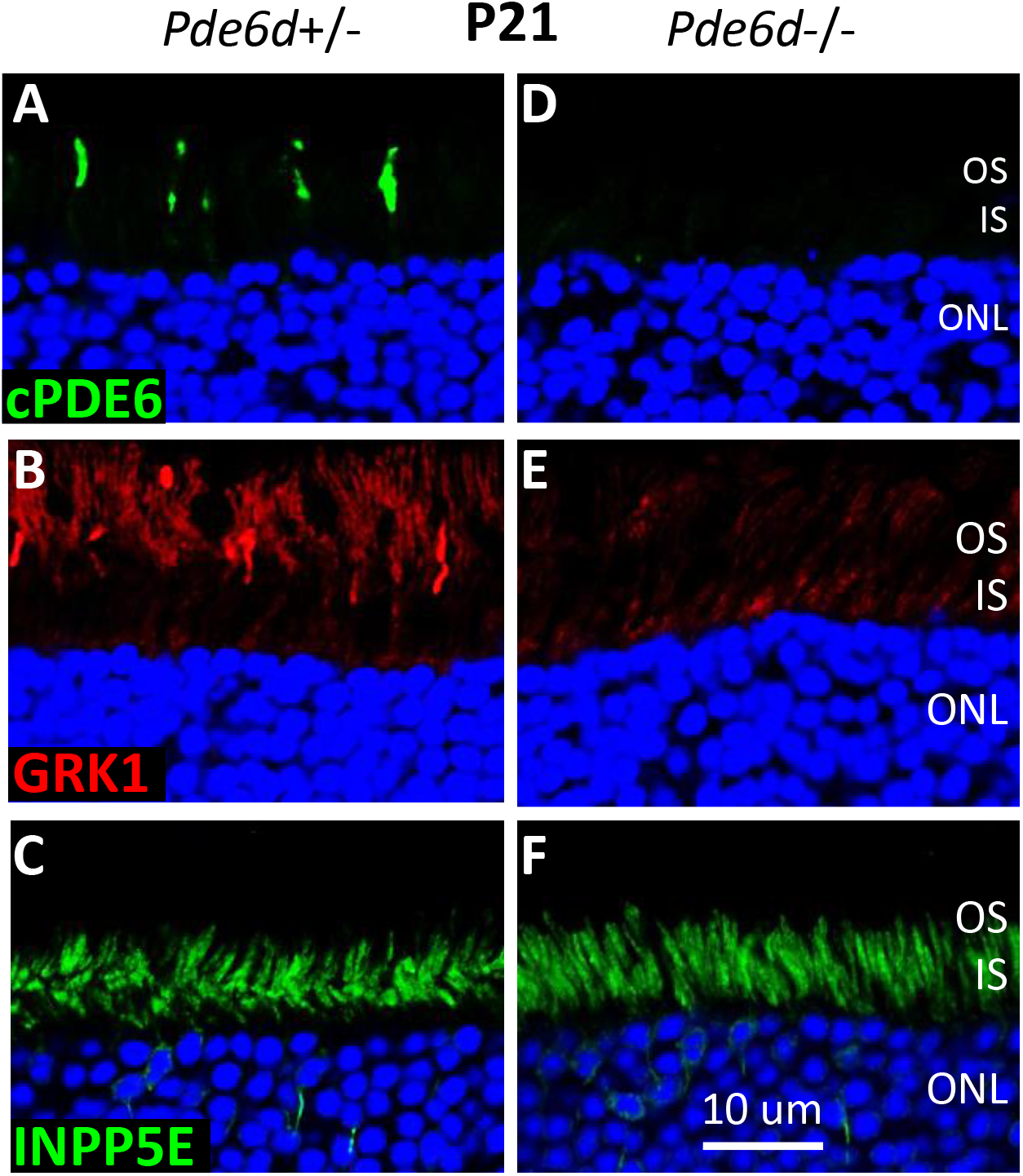
INPP5E localization in photoreceptors is independent of PDEδ. **A-F,** *Pde6d^-/-^* (A-C) and *Pde6d^-/-^* cryosections (D-F) were immunostained with anti-cPDE6 (A, D; green), anti-GRK1 (B, E; red) and anti-INPP5E (PT) (C, F; green) antibodies. Prenylated cone PDE6 and GRK1 require PDEδ (PDE6D) for OS localization. INPP5E inner segment localization is PDEδ-independent. Scale bar, 10 μm.

### Rhodopsin and PDE6 in ^ret^*Inpp5e^-/-^* retina

In WT and mutant retina at P6, OS begin to expand and rhodopsin is present in nascent WT OS and smaller mutant OS (**Fig. 6A, B;** P8 panels). At P10, mutant OS are shorter than the corresponding WT OS (**Fig. 6A, B;** P10 panel), rhodopsin begins to mislocalize and their connecting cilia (labeled with EGFP-CETN2) are located closer to ONL suggesting shrinking, degenerating IS. At P12, WT OS are maturing, but mutant OS are stunted and rhodopsin accumulates throughout the ONL (**Fig. 6A, B**, P12 panels). As judged by Western blotting (**Fig. 6C,** left panel), the level of rhodopsin in mutant rods is slightly reduced at P6 and about half at P10 and P12 (**Fig. 6C,** right panel).

**FIGURE 6.**
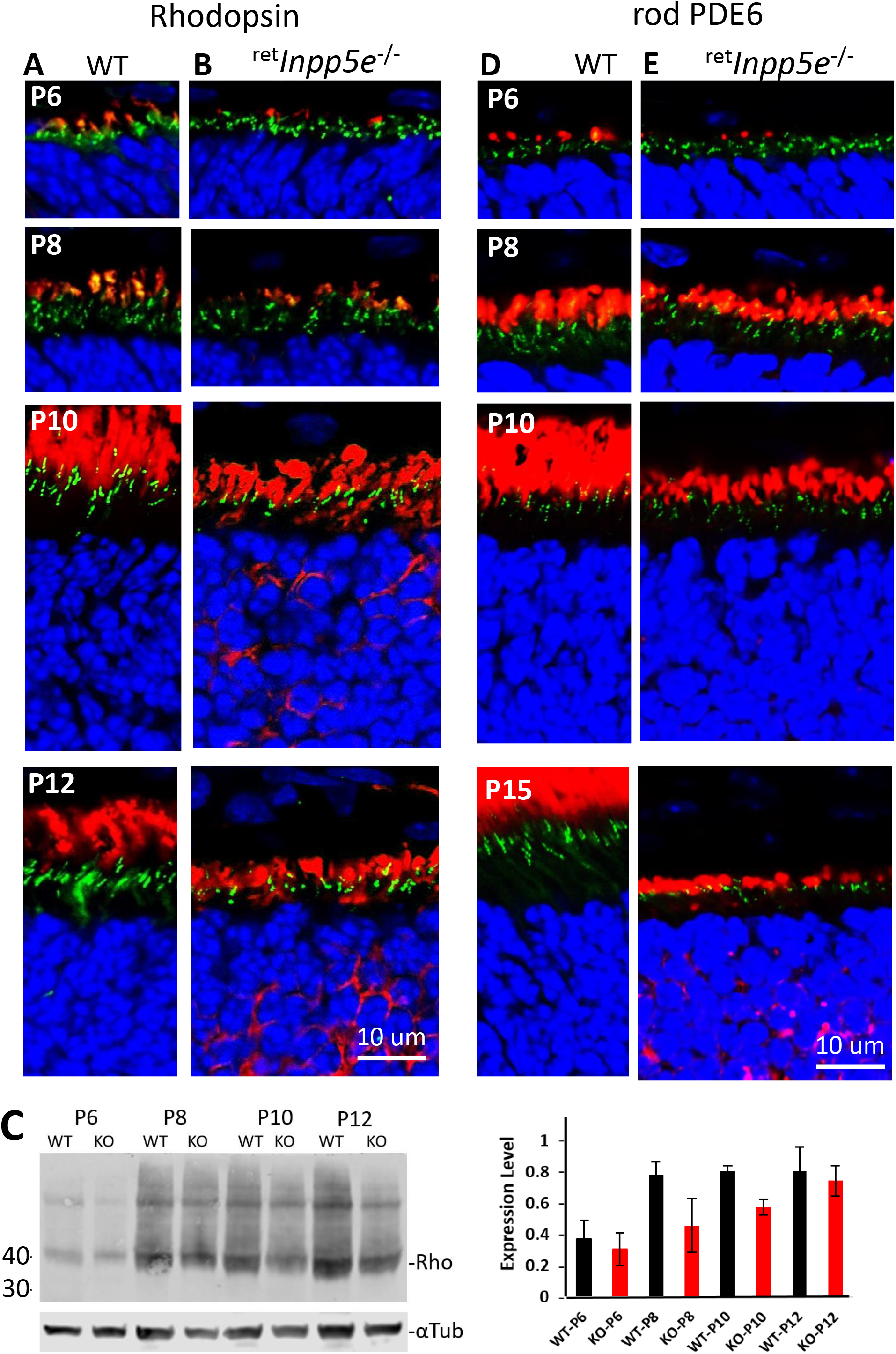
Immunolocalization of rhodopsin and rod PDE6 at P6 – P15. **A, B,** cryosections from WT (A) and ^ret^*Inpp5e^-/-^* (B) central retinas were probed with an anti-rhodopsin antibody (red) at P6, P8, P10 and P12. Transgenic EGFP-CETN2 (green) identifies centrioles and CC, DAPI (blue) stains nuclei. Starting at P10, photoreceptor OS do not expand and begin to degenerate. Scale bar, 10 um. **C,** Western blot of WT and knockout retina lysates at P6-P12. Densities of rhodopsin bands were normalized against the loading control α-tubulin and plotted on the graph on the right. **D, E**, cryosections from WT (D) and ^ret^*Inpp5e^-/-^* (E) central retinas were probed with an anti-PDE6 antibody (MOE) at P6, P8, P10, P15 and P21. Centrioles and connecting cilia are identified by transgenically expressed EGFP-CETN2. Nuclei are labeled with DAPI.

Prenylated peripheral OS proteins (PDE6, GRK1) are thought to traffic to the OS by diffusion, using PDEδ as a solubilization factor (27,28) and ARL3-GTP as a cargo dispensation factor (22,29,30). In WT and ^ret^*Inpp5e^-/-^* at P6, trace amounts of PDE6 are present in the OS (**Fig. 6D, E**). At P8 (just after the onset of degeneration), the level of OS PDE6 in ^ret^*Inpp5e^-/-^* rods is comparable to WT rods, but it is significantly reduced at P10 (**Fig. 6E**) consistent with the reduction of their OS size. These results suggest that ER-docking, for posttranslational modifications of PDE6 are not affected by INPP5E ablation.

### Mutant cones form spherical outer segments

Cone OS begin to form at P4 (31) and, in WT retina, can be detected by S-opsin immunofluorescence at P6 (**Fig. 7A**). At P8-P10, WT COS extend (**Fig. 7A**) while mutant COS form spherical structures (**Fig. 7B, 7B’**). At P12, WT COS are nearly mature while mutant COS remain spherical and increase in size. Similar spherical structures were seen when retinal sections were probed with antibodies against ML-opsin, cone PDE6, GRK1 and GUCY2E (GC1) (**Fig. 7C-F)**. Electron microscopy revealed that the mutant COS is filled with vesicles of various sizes and contains rudimentary fragments of axonemal extension. Yet, the CC appears to be normal in length (**Fig. 7G**). Large mitochondria (white arrow) at the distal IS confirm the cell is a cone.

**FIGURE 7.**
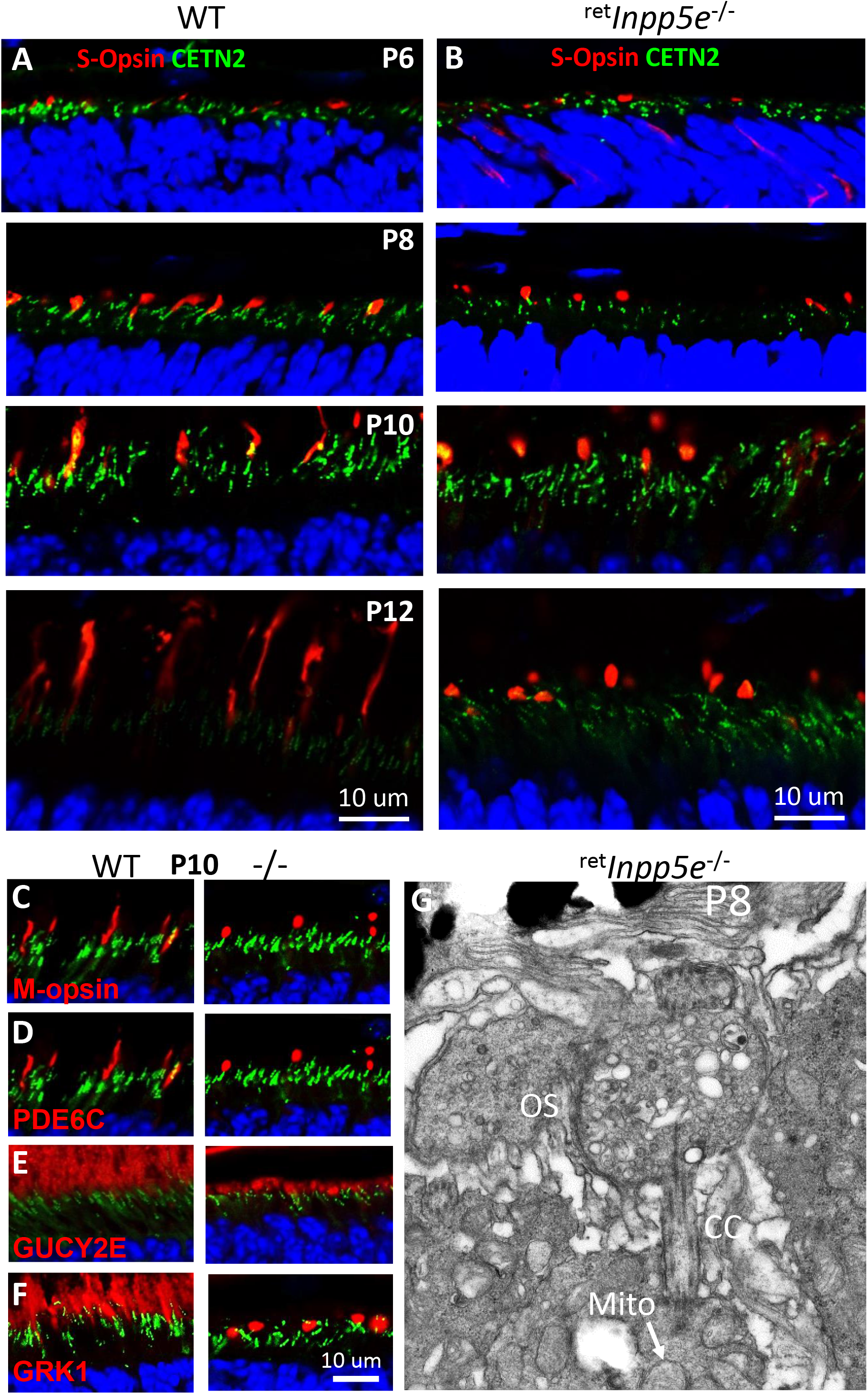
Mutant cones form spherical outer segments. **A, B**, S-opsin localization in WT (A) and ^ret^*Inpp5e^-/-^* (B) retinal cryosections at P6-P12. Cone outer segments are labeled with an anti-S-opsin antibody. CC are labeled by transgenically expressed EGFP-CETN2. In knockout mice, S-opsin is located in spherical cone OS at P8-P12. **C-F**, immunostaining of M-opsin (C), cone PDE6α’ (D), GC1 (E), and GRK1 (F) in WT (left panels) and mutant (right panels) retinas at P10. **G**, ultrastructural analysis of ^ret^*Inpp5e^-/-^* cone OS at P8. The OS appears as a spherical bag filled with vesicles and lacking OS axoneme extension and disc membranes. The identity of this cell as a cone is evident from the position if IS mitochondria (white arrow). See Ref. (53) for a representative image of a WT cone.

### Protein localization in mutant OS

Immunolocalization of OS resident proteins was performed at P10 using a battery of well-characterized antibodies (22,25) (**Fig. 8**). All investigated phototransduction proteins (GNAT1, PDE6 subunits, CNGA1/3 subunits) and structural (PRPH2, PROM1, CDHR1) proteins were present in mutant ROS (**Fig. 8**). ARL13b (known INPP5E interactant) and PRCD, involved in packaging membranes during disc morphogenesis (32), were located normally as well. An exception was the tubby-related protein TULP3 which was localized predominantly in the IS of WT rods (**Fig. 8I**, left panel), but was additionally present in at much greater amount in the mutant OS ((**Fig. 8I**, right panel).

**FIGURE 8.**
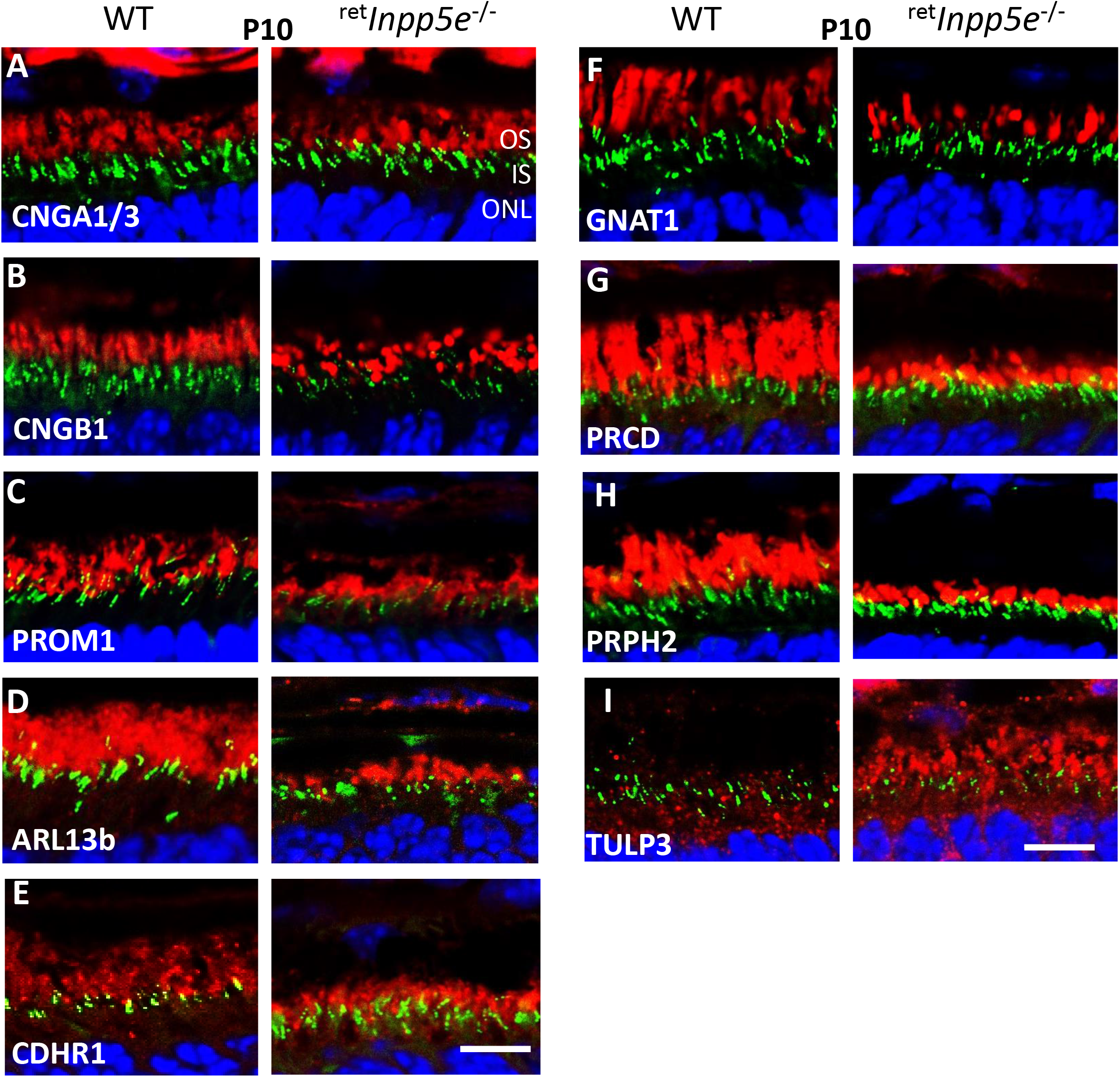
Survey of outer segment proteins’ localization in WT and ^ret^*Inpp5e^-/-^* retinas at P10. **A-I**, cryosections from WT (left) and knockout (right) retinas were probed with antibodies against CNGA1/3 (A), CNGB1 (B), PROM1 (C), ARL13B (D), CDHR1 (E), GNAT1 (F), PRCD (G), PRPH2 (*rds*) (H) and TULP3 (I). Scale bar, 10 μm.

### Ultrastructure of mutant rod outer segments

We next explored the ultrastructure of WT and mutant rod photoreceptors using transmission electron microscopy. At P6, rods produce primary cilium emanating from the basal body (**Fig. 9A**). The process of ciliogenesis is unaffected by the INPP5E knockout (**Fig 9C**). At P10, WT rods start forming OS containing regular disc structure (**Fig. 9B**). At this age, the CC structure of *^ret^Inpp5e^-/-^* rods remained normal and extended some axonemal microtubules into the proximal OS, but a normal -size axoneme was not generated. Importantly, the membrane structure emanating from the CC was highly disorganized and failed to form an ordered stack of discs (**Fig. 9D**).

**FIGURE 9.**
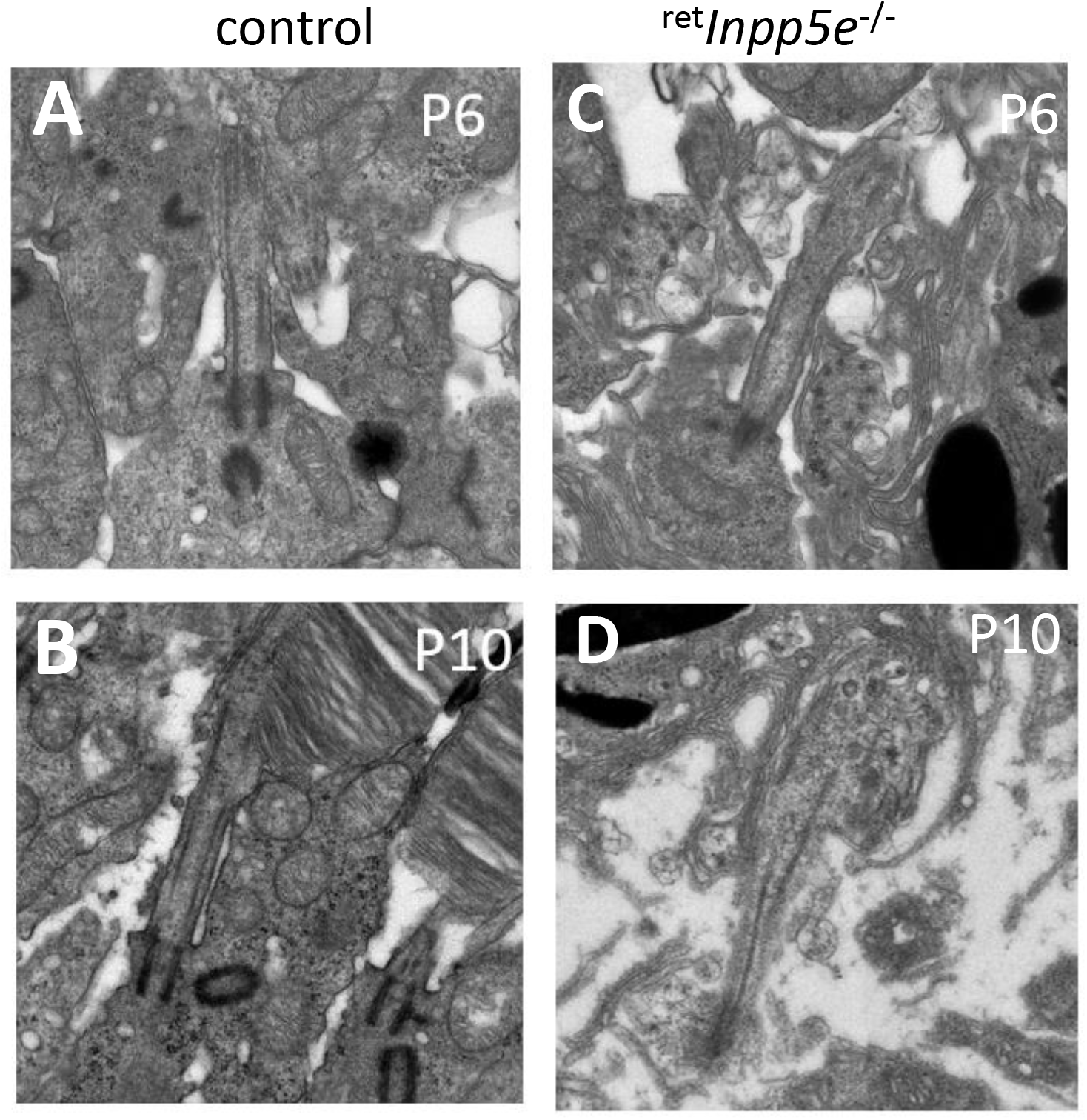
Comparison of ciliary and OS ultrastructure in WT and ^ret^*Inpp5e^-/-^* rods. **A, C**, representative images of cilia emanating from WT (A) and ^ret^*Inpp5e^-/-^* (C) rods at P6. **B, D,** representative images of WT (B) and ^ret^*Inpp5e^-/-^* (D) rods at P10 illustrating abnormal transition zone structure and rudimentary OS without discs in the knockout rod.

Immunolocalization conducted with retinal sections from WT and mutant mice (**Fig. 10A, B**) demonstrated correct positions of CEP290 (centrosomal protein 290), RPGR (Retinitis Pigmentosa GTPase regulator) and SPATA7 (spermatogenesis-associated protein 7), suggesting that mutant CC is intact. Overall, the *Inpp5e^-/-^* CCs appears to be indistinguishable from WT CC.

**FIGURE 10.**
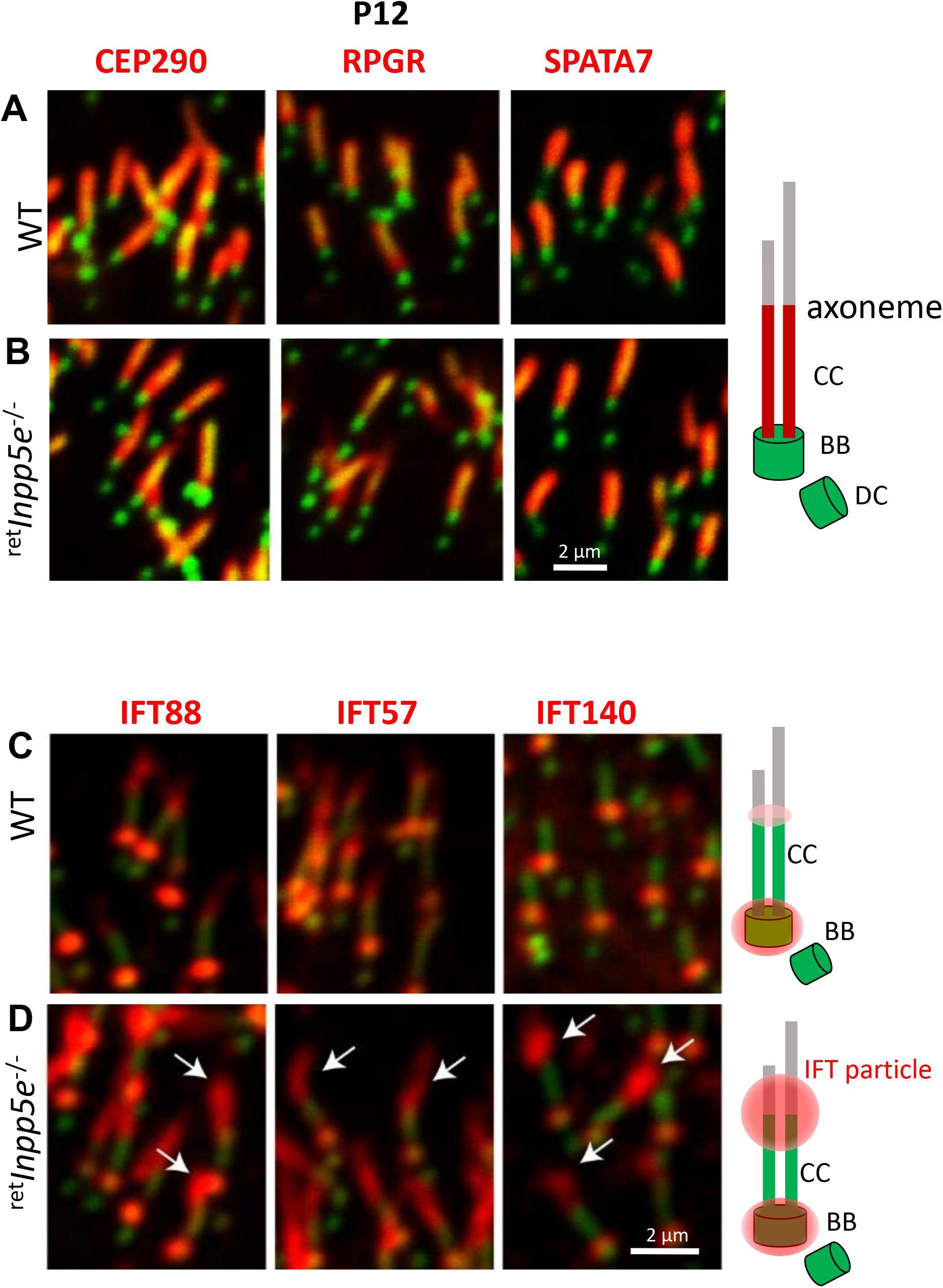
Mislocalization of intraflagellar transport proteins and normal distribution of ciliary transition zone proteins in ^ret^*Inpp5e^-/-^* rods. **A, B**, cryosections from WT (A) and ^ret^*Inpp5e^-/-^* (B) retinas collected at P12 were probed with antibodies against CEP290, RPGR and SPATA7 representing CC-specific proteins. Normal localization of these proteins suggests normal ultrastructure of the knockout CC. Scale bar, 2 μm. **C, D**, cryosections from WT (C) and ^ret^*Inpp5e^-/-^* (D) retinas collected at P12 were probed with antibodies against IFT88, IFT57 and IFT140. In WT photoreceptors, IFT particles are concentrated next to the basal body. Low levels of IFT88 and IFT57 are also observed in the proximal OS. In ^ret^*Inpp5e^-/-^* photoreceptors, a large fraction of IFT particles accumulates in the proximal OS (arrows). These changes are depicted in cartoons on the right.

### Impairment of IFT to form a normal axoneme

We next investigated the levels and positions of the IFT particles by immunostaining – IFT88, IFT57 and IFT140 (**Fig. 10C**). IFT88 and IFT57 are subunits of the IFT-B particle, whereas IFT140 is a subunit of the IFT-A particle complex. Normally, IFT particles are strongly associated with the basal body and proximal OS, and are essential for supporting anterograde and retrograde transport of the ciliary material (33-35). In P12 WT photoreceptors, IFT88 and IFT57 were strongly stained next to the basal body and weakly to the proximal OS, while IFT140 was seen only at basal body (**Fig. 10C**). In mutant photoreceptors IFT88, IFT57, and IFT140 massively accumulated at the CC/OS junction and in the proximal mutant OS (**Fig. 10D**, white arrows), suggesting impaired retrograde IFT.

As *Inpp5e^-/-^* OS development is severely impaired, we next explored how far the axoneme could extend into these altered OS structures. In WT control at P10, the axoneme visualized with an anti-acetylated α-tubulin (Ac-Tub) antibody (red) was extended from the CC (green) for approximately 4 μm (**Fig. 11A, C**). In mutant rods the axoneme was stunted and never exceeded 1 μm (**Fig. 11B, C**), consistent with TEM results (**Fig. 9D**).

**FIGURE 11.**
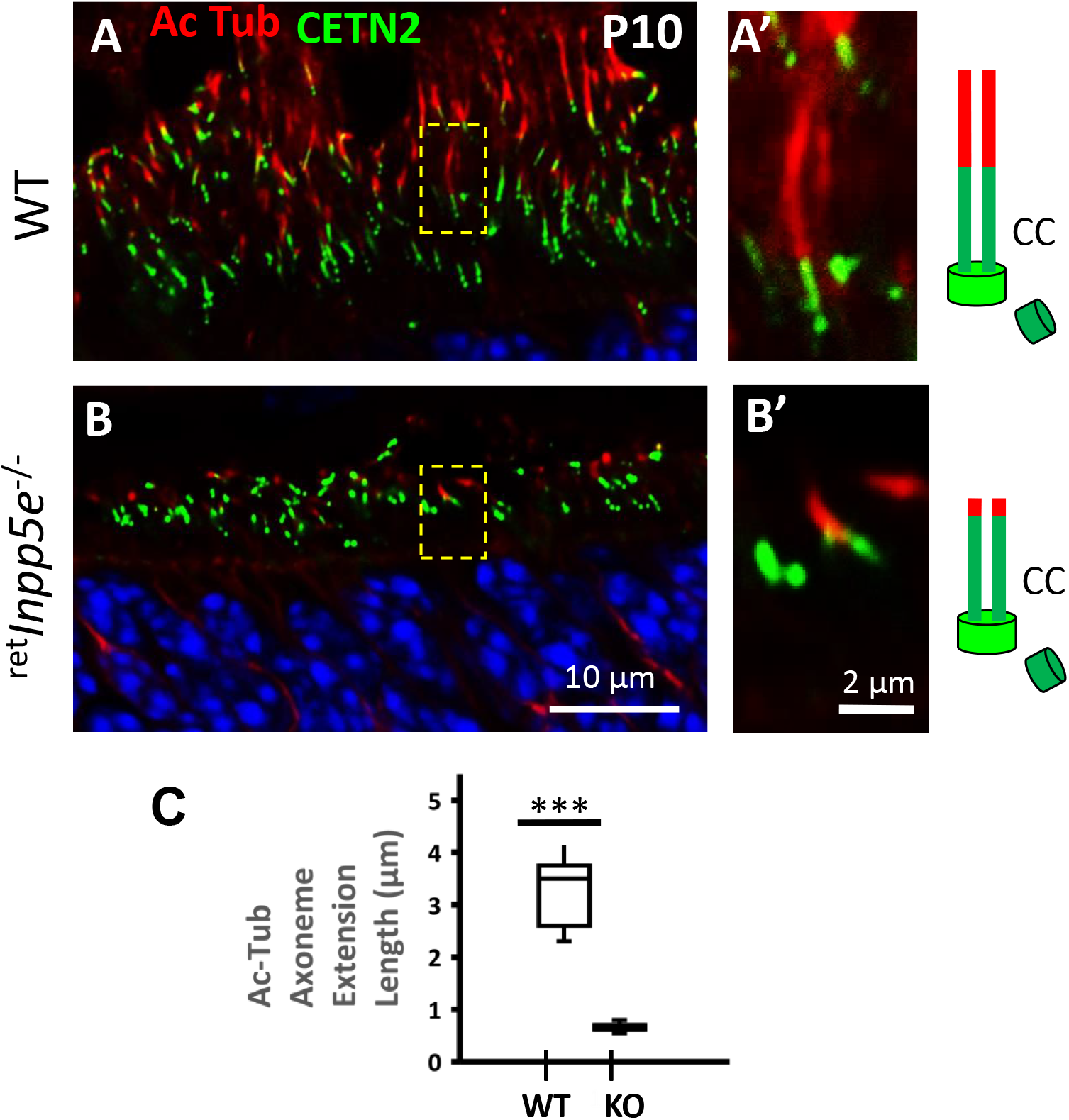
Stunted axoneme in developing ^ret^*Inpp5e^-/-^* photoreceptors. **A, B,** ^ret^*Inpp5e^+/-^* (A) and ^ret^*Inpp5e^-/-^* (B) retinal cryosections were stained with an antibody against acetylated α-tubulin (Ac-Tub). Panels A’ and B’ are enlargements. Cartoons illustrating basal bodies and CC (green) and axonemes (red) are shown on the right. WT Ac-Tub positive axonemes range between 2 and 4 μm in length, whereas knockout axonemes are severely stunted. **C**, quantification of the axoneme length. Student’s *t*-test analysis of two datasets was performed assuming equal variance. P= 3.22946E-10. n = 3.

## DISCUSSION

In this study, we demonstrated that INPP5E localizes to the photoreceptor inner segments and explored the consequences of its retina-specific deletion. INPP5E is a farnesylated phosphatidylinositol polyphosphate 5’-phosphatase regulating the level of phosphoinositides in the primary cilium (5,6). Its ciliary targeting in IMCD3 cells was shown to be PDEδ/ARL3-dependent. Absence of PDEδ or inhibiting PDEδ activity prevented INPP5E delivery to the cilium of MEFs (17).

Using immunostaining with two different antibodies and serial tangential sectioning with immunoblotting (**Fig. 3**), we show that INPP5E is a predominantly IS protein, but is also present in the ONL (**Fig. 4A, C, E, G**) and the CC (**Fig. 4J, L**). INPP5E is posttranslationally farnesylated requiring docking to the ER for CAAX box processing (removal of –AAX from the CAAX box and carboxymethylation) (36). INPP5E is retained in part in the ONL (**Fig. 4)**, presumably docked to the perinuclear ER. However, the bulk of INPP5E is present in the IS, independent of PDEδ (**Fig. 4F**), or ARL3-GTP which serves as a cargo dispensation factor for OS prenylated proteins (22). Why INPP5E is retained at IS membranes and why it is excluded from the OS is unclear.

^ret^*Inpp5e^-/-^* OSs are stunted beginning at P8 and neither reach normal length nor elaborate normal ultrastructure. At P10, mutant photoreceptors start to degenerate, rhodopsin mislocalizes in the ONL and the ONL thickness is reduced (**Fig. 2**). The amount of rhodopsin in mutant OS appears reduced starting at P8. (**Fig. 6C**). The mislocalization of rhodopsin beyond P10 and PDE6 beyond P12 (**Fig. 6B**) are likely secondary effects of failed OS extension. Rod and cone ^ret^*Inpp5e^-/-^* OS form an apparently normal length CC (**Fig. 9**) but do not extend an axoneme into the OS and do not form discs (**Fig. 10**), suggesting an involvement of phosphoinositides in axoneme elongation and disc morphogenesis. Presence of CC and absence of OS axoneme extensions and discs has been observed in a number of animal models lacking OS proteins, including rhodopsin knockouts (37,38), *rd1* mouse (PDE6b null allele) (39) and PRPH2 (*rds*) (40) knockouts. Presence of CC and absence of OS also was observed in deletions of a number of ciliary proteins including NPHP5 (41). NPHP1 (42), AHI1 (jouberin) (43), lebercilin (LC5) (44), demonstrating that axoneme extension and disc morphogenesis is complex process requiring a large number of components.

One possible explanation of the INPP5E knockout phenotype is that trafficking of key OS proteins is impaired in the absence of INPP5E. However, our analysis of intracellular localization of several OS-resident proteins by confocal microscopy showed that all of them were normally trafficked to the mutant OS (**Fig. 8**). This shows that INPP5E is dispensable for ciliary targeting of key OS proteins.

PIPs have important functions in membrane signaling and cytoskeleton dynamics (4). The major role of INPP5E in primary cilia is to create a PI4P-rich ciliary membrane environment by catalyzing the conversion of PIP2 to PI4P, which is essential to regulate the ciliary localization of receptor proteins of the SHH signaling pathway. In photoreceptors, the concentration of PIPs in rod IS and OS is very low at roughly 0.04 mol% (45). Recent measurements with fluorescent phosphoinositide sensors revealed that the majority of PI4P and PI(4,5)P2 rods is present in the IS and cytoplasm surrounding the nuclei (46); the OS also contains a smaller amount of PI4P but no detectable PIP2. The latter is consistent with a direct measurement showing the presence of trace amount of PI4P but not PI(4,5)P2 in isolated bovine outer segments (47)

Deletion of INPP5E is expected to decrease PI4P and to increase the concentrations of PIP2 in all cellular compartments. The increase of TULP3 in P10 KO (**Fig. 8I**) is consistent with this prediction. Tubby family proteins (Tubby, Tulp1-4) are known PIP2 interactors (48) required for ciliary entry of certain GPCRs by bridging the IFT-A complex with PIP2 (49,50). The increase of TULP3 may explain the accumulation of IFT proteins at the OS base (**Fig. 10D**) as TULP3 binds to IFTA (49,50)). Accumulation of PIP2, TULP3 and IFT particles was also observed in *Inpp5e^-/-^* primary cilia of neural stem cell cultures (5).

The accumulation of IFT particles at OS base suggests a defective retrograde IFT. Retrograde IFT is essential for the OS formation and extension, as morpholino knockdowns of the dync2h1, dync2li1 or dync2i1 subunits of retrograde IFT motor cytoplasmic dynein-2 lead to dramatic OS shortening or complete lack of OS (51). Future studies will be directed to elucidating the role of PIPs and IFT in defects in disc morphogenesis and axoneme extension caused by the INPP5E deficiency.

## EXPERIMENTAL PROCEDURES

### Animals

All procedures were approved by the University of Utah Institutional Animal Care and Use Committee and were conducted in compliance with the NIH Guide for Care and Use of Laboratory Animals. Floxed *Inpp5e* mice (*Inpp5e^F/F^*), provided by Dr. Christina Mitchell, were maintained in a 12:12 h dark-light cycle. A transgenic mouse expressing EGFP-CETN2 fusion protein (JAX stock # 008234) was used to identify centrioles/transition zones (52). Three month-old Long-Evans rats used for tangential retinal sectioning were obtained from Charles River.

### *Generation of ^ret^Inpp5e^-/-^* conditional *knockout mice*

*Inpp5e^F/F^* mice (12) were mated with Six3-Cre transgenic mice (24) to delete exons 2-5, thereby generating retina-specific *Inpp5e* knockouts (^ret^*Inpp5e^-/-^*) (**Fig. 1**). Knockout mice were viable but had a reduced litter size (4-5 pups). *Inpp5e* floxed allele PCR reactions were performed on genomic DNA (13) using *Inpp5e* WT forward primer 5’-GAGAAGCTGATAGATGGCTAGG and *Inpp5e* WT reverse primer 5’-AACCAGAAGACCTCATCAAACC and EconoTaq PCR according to manufactures specifications (Lucigen corporation, WI) (**Fig. 1D**). Homologous recombination in retina was verified with INPP5E-KO-F primer 5’-CAGAATGCATAGCTCTCTGGGCAAC and INPP5E WT-R primer 5’-GTAGTGACATCCCCTGGGCACGTG (amplicon size 450 bp) using retina DNA as a template. The *Inpp5e* floxed allele amplicon is 429 bp, the *Inpp5e* wild-type allele is 300 bp, and the knockout amplicon is 450 bp. Six3-Cre mice were genotyped with Cre-specific primer set Six3Cre159 forward 5’-TCGATGCAACGAGTGATGAG and Six3Cre160 reverse primer 5’-TTCGGCTATACGTAACAGGG (amplicon size 500 bp). The Egfp-Cetn2+ transgene was identified with the primer set, Egfp-Cetn2+-F 5’-TGAACGAAATCTTCCCAGTTTCA and Egfp-Cetn2+-R 5’-ACTTCAAGATCCGCCACAACAT (amplicon size 600 bp). PCR amplicons were separated using 1.5% agarose gel electrophoresis in the presence of ethidium bromide and visualized via transillumination. Absence of the *rd8* and *rd1* mutation was confirmed by PCR (53).

### Immunoblotting

For western blotting, two retinas from one mouse were homogenized in 100 μl of 50 mM Tris-HCl (pH 8), 100 mM NaCl, 10 mM EDTA and 0.2 % Triton X-100, and 2 μl of 100 mM PMSF and 2μl protease inhibitor. Samples were then sonicated for 2 x 20 pulses at an intensity of 30% and spun at 15,000 rpm for 20 min. The protein concentration was determined by Bradford assay. Proteins of retina lysates were separated by 10% SDS-PAGE and transferred to a nitrocellulose membrane (54). The resulting membrane was sequentially subjected to blocking for 1 h, primary antibody incubation overnight at 4°C and secondary antibody incubation for 1 h. Primary antibodies were diluted 1:500 for anti-INPP5E (ProteinTech), 1:5000 for anti-Rho (4D2). Secondary antibodies were iR680 goat-anti-mouse (1:5000) and iR800 goat-anti-rabbit (1:3000) (Odyssey); images were acquired using an Odyssey scanner. The intensities of protein bands were measured using Image J and normalized to control bands.

### Confocal immunohistochemistry

Animals were dark-adapted overnight and sacrificed in ambient light. For conventional fixation (55), P6-P21 eyes from control and mutant mice were immersion-fixed using freshly prepared 4% paraformaldehyde in 0.1 m phosphate buffer, pH 7.4, for 2 h on ice. Eyecups were then moved to 15% sucrose in phosphate buffer for 1 h and then to 30% sucrose overnight for cryoprotection.

For low fixation protocol (CO INPP5E antibody and MAK CC marker do not work with standard fixation methods), mouse eyes were rinsed in 1xPBS followed by embedding in OCT. Blocks were immediately frozen at −80°C, and sectioned at 14um. Slides were removed from freezer and warmed no longer than 5 min. 1% PFA (made by diluting 4% PFA in PBS) was applied to the slide for 2 min. slides are washed 5 min in 1xPBS. Sections were blocked using 10% normal goat serum in 0.1 m phosphate buffer–0.1% Triton X-100 (PBT) for 1 h and incubated with primary antibody overnight in a rotating humidified chamber at 4°C.

Cryosections were incubated with the following polyclonal primary antibodies: rod anti-transducin-α (1:500, Santa Cruz). Anti-M/L-opsin (1:500, Chemicon). Anti-S-opsin (1:500, Chemicon). MOE (anti-rod PDE6, 1:500, Cytosignal). Anti-cone PDE6 (1:500, gift of Dr. Tiansen Li, NEI). PRCD, Vadim Arhavsky lab (32). PROM1 (ProteinTech) (1: 400). CNGA1/3 (1:1000. NeuroMab, UCDavis). CDHR1 (1:500, Jun Yang, University of Utah). SPATA7 (1:100, Rui Chen, Baylor College of Medicine). Rat CEP290 (1:300, Anand Swaroop, NEI). IFT88, IFT57 and IFT140 (1:500 Greg Pazour). TULP3 (1:200, ProteinTech). Monoclonal antibodies included: Ac-Tub (1: 1000, Sigma). G8 (1:500, anti-GRK1, Santa Cruz). PRPH2 (2B6) (1:25, Bob Molday, University of British Columbia). GC1 (IS4) (1:1000, Kris Palczewski, UC Irvine). Anti-ARL13b (1:200, NeuroMab). Cy3- or Alexa488-conjugated goat-anti-rabbit and goat-anti-mouse secondary antibodies were diluted 1:1000 in blocking solution (2% BSA, 0.1M phosphate buffer, pH 7.4, containing 0.1% Triton X-100). Images were acquired using a Zeiss LSM800 confocal microscope.

### Retinal serial sectioning with western blotting

Experiments were performed as described in (56,57). Rats were sacrificed, and eyes were enucleated and dissected in ice-cold Ringer’s solution. A retina punch (3 mm diameter) was cut from the eyecup with a surgical trephine positioned adjacent next to the optic disc, transferred onto PVDF membrane with the photoreceptor layer facing up, flat mounted between two glass slides separated by plastic spacers (ca. 240 μm) and frozen on dry ice. The retina surface was aligned with the cutting plane of a cryostat and uneven edges were trimmed away. Progressive 5-μm tangential sections were collected, subjected to SDS-PAGE and probed with antibodies against INPP5E (ProteinTech) and rhodopsin (4D2).

### Measurement of ONL thickness

Mouse eye cups, with the anterior segments and lens removed, were fixed overnight in 4% PFA in PBS overnight. Eyes from both control and mutant mice were then immersed in 15% sucrose in phosphate buffer for 1 h and then to 30% sucrose overnight for cryoprotection. Twelve-micron transverse-sections were stained with DAPI and the thicknesses of the ONL layers were measured along the retinal vertical meridian at approximately 500um apart on each side of the optic nerve head.

### Electroretinography (ERG)

Scotopic and photopic electroretinogram (ERG) responses were recorded from P15 WT, ^ret^*Inpp5e*^+/-^ and ^ret^*Inpp5e^-/-^* mice using a UTAS BigShot Ganzfeld system (LKC Technologies, Gaithersburg, MD, USA). ERGs were measured as described (22,58).

### Transmission Electron Microscopy

Isolated mouse eyecups at ages of P8 and P10 were fixed by immersion in fixative (2% glutaraldehyde-1% paraformaldehyde in 0.1M cacodylate buffer, pH 7.4) at 4°C overnight (32, 46). The eyecups were postfixed with 1% osmium tetroxide in 0.1M cacodylate for 1 h, buffer-rinsed, stained *en bloc* with uranyl acetate, and subsequently dehydrated in an ascending series of methanol solutions. Eyecups were embedded in Epon resin (Ted Pella, Inc., Redding, CA) for sectioning. 1 μm plastic sections were cut to face and orient photoreceptors near the optic nerve. Retina ultrathin (60 nm) sections were cut and transferred onto slot grids with carbon-coated Formvar film (Electron Microscopy Sciences, Hatfield, PA) and poststained with uranyl acetate followed by lead citrate. Transmission electron microscopy was performed at 75 kV using a JOEL electron microscope.

### Statistics

SigmaPlot12 was used for statistical analysis using student *t*-test and the level of statistical significance was set *P* = 0.05.

## Acknowledgement

We thank the following investigators for providing antibodies: Dr. Robert Molday (University of British Columbia-Vancouver) for rhodopsin (1D4), Rom1 and peripherin2 antibodies; Dr. Kris Palczewski (UC-Irvine) for GC1 (IS4) antibody; Dr. Wieland Huttner (MPI Dresden/Germany) for prominin1 antibody; and Dr. Greg Pazour (University of Massachusetts Medical School) for IFT88, IFT157 and IFT140 antibodies.

## Author contributions

AS and WB designed the study; AS, GY, CDG JMF and MAC performed research; AS, GY, JMF, VYA and WB analyzed data; WB wrote the manuscript; JMF, GY and VYA edited the manuscript.

## Footnotes

This work was supported by NIH grants EY08123, EY019298 (WB); EY012859 (VYA); EY005722 (VYA); EY014800-039003 (NEI core grant). 5T32 EY024234 (NEI training grant); by unrestricted grants to the University of Utah Department of Ophthalmology from Research to Prevent Blindness (RPB; New York) and by a grant from the Retina Research Foundation Houston (Alice McPherson, MD). WB and VYA are recipients of the RPB Senior Investigator award and the RPB Nelson Trust award. AS is a fellow on the NEI training grant.

